# Activation of the aryl hydrocarbon receptor inhibits neuropilin-1 upregulation on IL-2 responding CD4^+^ T cells

**DOI:** 10.1101/2023.09.25.559429

**Authors:** Simone Sandoval, Keegan Malany, Krista Thongphanh, Clarisa A. Martinez, Michael L. Goodson, Felipe Da Costa Souza, Lo-Wei Lin, Jamie Pennington, Pamela J. Lein, Nancy I. Kerkvliet, Allison K. Ehrlich

## Abstract

Neuropilin-1 (Nrp1), a transmembrane protein expressed on CD4^+^ T cells, is mostly studied in the context of regulatory T cell (Treg) function. More recently, there is increasing evidence that Nrp1 is also highly expressed on activated effector T cells and that increases in these Nrp1-expressing CD4^+^ T cells correspond with immunopathology across several T cell-dependent disease models. Thus, Nrp1 may be implicated in the identification and function of immunopathologic T cells. Nrp1 downregulation in CD4^+^ T cells is one of the strongest transcriptional changes in response to immunoregulatory compounds that act though the aryl hydrocarbon receptor (AhR), a ligand-activated transcription factor. To better understand the link between AhR and Nrp1 expression on CD4^+^ T cells, Nrp1 expression was assessed *in vivo* and *in vitro* following AhR ligand treatment. In the current study, we identified that the percentage of Nrp1 expressing CD4^+^ T cells increases over the course of activation and proliferation *in vivo*. The actively dividing Nrp1^+^Foxp3^-^ cells express the classic effector phenotype of CD44^hi^CD45RB^lo^, and the increase in Nrp1^+^Foxp3^-^ cells is prevented by AhR activation. In contrast, Nrp1 expression is not modulated by AhR activation in non-proliferating CD4^+^ T cells. The downregulation of Nrp1 on CD4^+^ T cells was recapitulated *in vitro* in cells isolated from C57BL/6 and NOD (non-obese diabetic) mice. CD4^+^Foxp3^-^ cells expressing CD25, stimulated with IL-2, or differentiated into Th1 cells, were particularly sensitive to AhR-mediated inhibition of Nrp1 upregulation. IL-2 was necessary for AhR-dependent downregulation of Nrp1 expression both *in vitro* and *in vivo*. Collectively, the data demonstrate that Nrp1 is a CD4^+^ T cell activation marker and that regulation of Nrp1 could be a previously undescribed mechanism by which AhR ligands modulate effector CD4^+^ T cell responses.

## 1 Introduction

The aryl hydrocarbon receptor (AhR) is an attractive therapeutic target for regulating proinflammatory immune responses (1). As a highly conserved ligand-activated transcription factor, AhR alters the expression of genes involved in the activation, differentiation, and survival of immune cells (2, 3). In particular, newly activated CD4^+^ T cells are primary targets of AhR-mediated immune regulation (4). Because of its immunomodulatory properties, AhR activation has been studied for the treatment of several immune-mediated diseases including type 1 diabetes (5), graft-versus-host (GVH) disease (6), inflammatory bowel disease (7), and experimental autoimmune encephalomyelitis (8). These studies consistently demonstrate that AhR ligands promote Foxp3^+^ Treg and Tr1 differentiation, although it is less understood how AhR signaling directly modulates proinflammatory CD4^+^ T cell subsets.

Global gene expression analysis of newly activated CD4^+^ T cells demonstrated that two high affinity AhR ligands (the metabolically resistant ligand 2,3,7,8-tetrachlorodibenzo-p-dioxin; TCDD) and rapidly metabolized ligand, 11-chloro-7H-benzimidazo[2,1-a]benzo[de]iso-quinolin-7-one (11-Cl-BBQ); 11-Cl-BBQ), induce an overlapping gene signature; an increase in AhR-responsive genes as well as those involved in T cell activation, immunoregulation, and migration (9). These changes are accompanied by a decrease in genes involved in cell cycling and adhesion. The gene encoding neuropilin-1 (Nrp1) was among the most highly downregulated following AhR activation (9). Nrp1 is a transmembrane glycoprotein with broad ligand specificity which includes secreted class III semaphorins, vascular endothelial growth factor (VEGF), TGF-β, TGF-βR1/II/III, as well as Nrp1 itself (10, 11). Consistent with this diverse functionality, Nrp1 is expressed on several cell types including neurons, keratinocytes, osteoblasts, endothelial cells, and immune cells (10-13). Initially, Nrp1 was recognized for its role in normal embryonic development, axon guidance, and vasculature formation (10, 13). However, since its first discovery, Nrp1 has been further implicated in the function of dendritic cells, macrophages, and T cells, where it is involved in establishing the immunological synapse and the generation of the primary immune response (12, 14). Within the context of CD4^+^ T cells, Nrp1 has been mostly studied as a marker for thymically derived Tregs as its expression is positively correlated with Foxp3 during Treg development and contributes to their function and stability (15-17).

Although mostly studied on Tregs, several reports have suggested that Nrp1 is also expressed on activated proinflammatory CD4^+^ T cells (18, 19). We previously found that Nrp1^+^ Foxp3^-^ cells positively correlate with insulitis in non-obese diabetic mice (NOD) (18) and with cytotoxic T lymphocytes (CTL) in a graft-versus-host model (19). In both models, 11-Cl-BBQ and TCDD prevent disease development in an AhR-dependent manner (18, 20) and lead to reduced Nrp1 expression on CD4^+^Foxp3^-^ cells. This finding is not entirely surprising given the role of Nrp1 in the generation of the primary T cell response. Additionally, Abberger, et al. (21) recently reported that Nrp1 expression is induced upon activation of CD4^+^ Foxp3^-^ T cells, and these cells express a highly activated phenotype including elevated proliferative capacity and an increase in CD44, CD25, and proinflammatory cytokines. In the present studies, we characterize Nrp1 expression on CD4^+^ T cell subsets *in vivo* and *in vitro* and propose AhR-IL-2-Nrp1 signaling as an additional mechanism by which drugs that activate AhR lead to the regulation of proinflammatory CD4^+^ T cell responses.

## 2 Materials and Methods

### Animals

NOD, C57BL/6J (B6; H-2^b/b^) and B6D2F1 (F1; H-2^b/d^) mice were originally purchased from The Jackson Laboratory and bred and maintained in a specific pathogen-free animal facility at University of California, Davis or at Oregon State University. NOD.AhR^−/−^ mice were generated, as previously published (18). Timed pregnant Sprague-Dawley rats (RRID: MGI_5651135) were obtained from Charles River Laboratory received at the University of California, Davis at least 16 d post-conception (E16), where they were individually housed at constant temperature (∼22°C), with a 12-h light-dark cycle until delivery. All animal procedures were carried out following protocols approved by the Institutional Animal Care and Use Committee at Oregon State University and University of California, Davis.

### Donor Cell Transfer

Donor cells were pooled from the spleen and peripheral lymph nodes of B6 mice, and 3-4 × 10^7^ donor T cells were transferred through tail vein injection into F1 host mice as we previously described (9, 19). Prior to donor cell injection, cells were labeled with 5 μM carboxyfluorescein succinimidyl ester (CFSE; Life Technologies; Cat # C34554). For Day 2 studies, donor cells were identified based on CFSE staining. For studies at later time points, donor cells were also differentiated from host cells by antibodies targeting H-2D^d^.

### AhR ligand treatments

For *in vivo* studies, TCDD (Cambridge Isotope Laboratories; item # ED-901-C) was dissolved in anisole, further diluted in peanut oil and administered at 15 μg/kg TCDD. 11-Cl-BBQ (ChemBridge) was dissolved in dimethyl sulfoxide (DMSO), further diluted in peanut oil, and mice were administered 10 mg/kg. The vehicles used for both TCDD and 11-Cl-BBQ were a 0.15% anisole solution and a 1% DMSO solution made in peanut oil, respectively. The route of administration was intraperitoneal injection daily. For *in vitro* studies, 11-Cl-BBQ was dissolved in DMSO and further diluted in supplemented culture medium.

### *In vitro* CD4^+^ T cell stimulation

Single cell suspensions of splenocytes were prepared and CD4^+^ T cells were isolated using the EasySep™ Mouse CD4^+^ T Cell Isolation Kit (StemCell Technologies; Cat # 19852A). CD4^+^ T cells were plated in a round bottom 96-well plate at 2 × 10^5^ cells/well and activated using Mouse T Activator CD3/CD28 Dynabeads™ (Gibco; Cat # 11456D). 11-Cl-BBQ was added at a concentration of 100nM. CD4^+^ T cells were differentiated according to Table 1 as previously described (22). After 4 days of incubation, CD3/CD28 Dynabeads were removed through magnetic separation. Cells were collected for real-time PCR or flow cytometry.

**Table 1:**
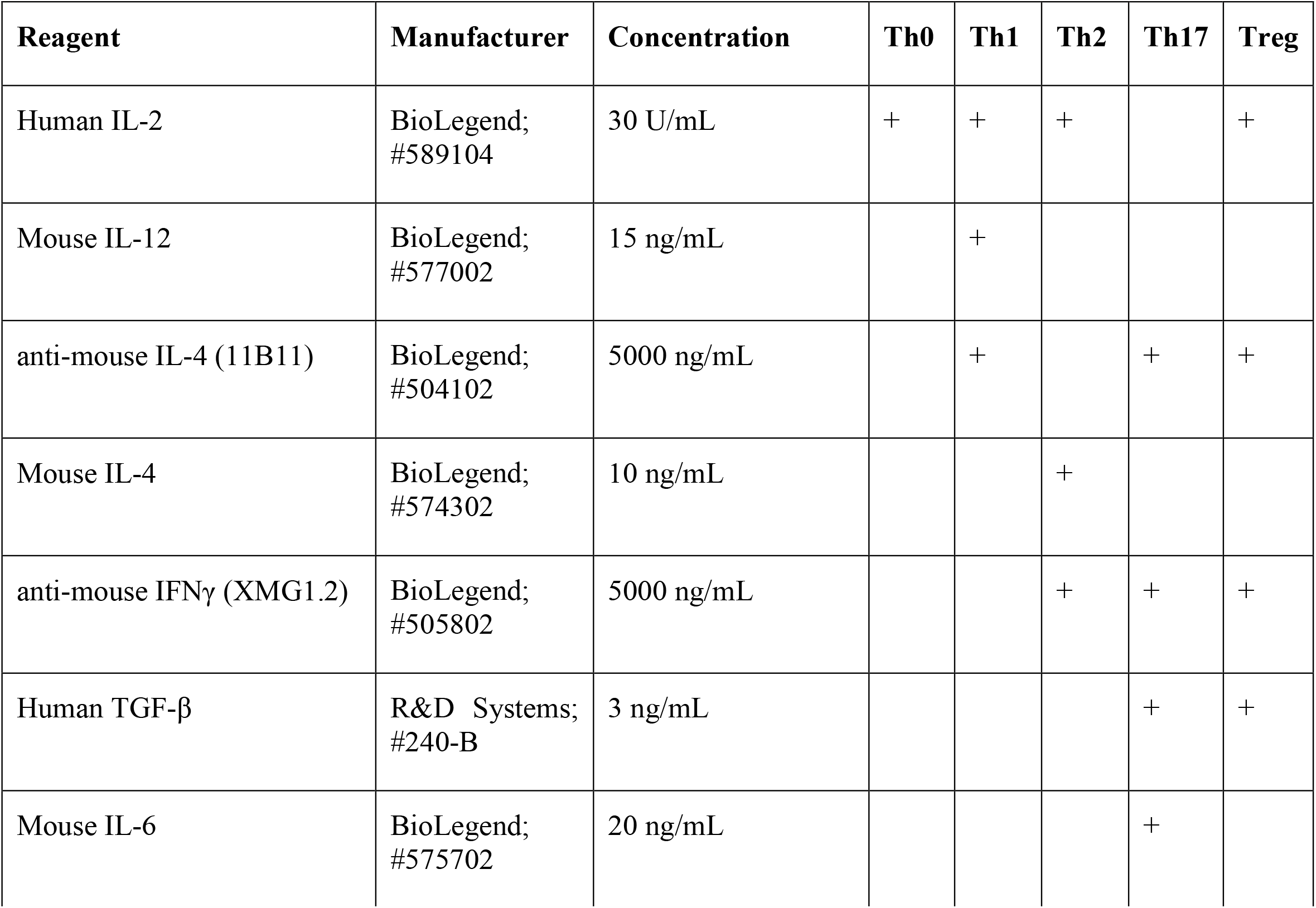
Conditions to differentiate into CD4^+^ T cell subsets.

#### *In vitro* IL-2 neutralization

Th1 polarizing reagents were added to CD4^+^ T cells without IL-2. Autocrine production of IL-2 was neutralized by the addition of an anti-IL2 antibody (*InVivo*MAb anti-mouse IL-2; clone JES6-1a12; BioXcell; Cat # BE0043-1) at 10 ng/ml, 100 ng/ml and 1000 ng/ml. 11-Cl-BBQ was added at a concentration of 100 nM. Following incubation at 37°C for 4 days, CD3/CD28 Dynabeads were removed through magnetic separation and cells were collected for flow cytometry.

#### *In vivo* IL-2 neutralization

At the time of donor cells transfer, 0.5mg of anti-IL2 antibody mixture or isotype control (IgG2a) was injected i.p. into F1 host mice. The anti-IL2 contained a 1:1 mixture of S4B6 (BioXcell; Cat # BE0043-1) and JES6-1a12 (BioXcell; Cat # BE0043) antibodies.

### Flow cytometry

Cells were stained with fixable viability dye (Fixable Viability Dye eFluor 780; eBioscience; Cat # 65-0865-14). Fc receptors were blocked with anti-CD16/CD32 (Anti-mouse CD16/CD32; clone 93; eBioscience; Cat # 14016185) and the cells were stained with antibodies specific for the following proteins: CD4 (PE Rat Anti-Mouse CD4; Clone RM4-5; BD Biosciences; Cat # 553049), H-2D^d^ (Anti-mouse H-2D^d^ antibody; clone 34-2-12; Biolegend; Cat #110606), CD25 (BV510 Rat Anti-Mouse CD25; clone PC61; BD Biosciences; Cat # 563037), CD29 (APC-Efluor 780 Anti-mouse CD29 antibody; clone eBioHMb1-1; eBiosciences; Cat # 47029182), CD69 (APC-Efluor780 Anti-mouse CD69 antibody; clone H1.2F3; eBiosciences; Cat # 47069182) and Nrp1 (PE/Cyanine7 anti-mouse CD304 (Neuropilin-1); clone 3E12; BioLegend; Cat #145211). For intracellular staining, cells were fixed and permeabilized using the Foxp3 Fixation/Permeabilization buffer kit (eBioscience; Cat # 00552300) and stained with antibodies specific for the following proteins: Foxp3 (PE-Cyanine5 FOXP3 Monoclonal; clone FJK-16s; eBiosciences; Cat # 15577382). For in vitro studies, cells were stained with fixable viability dye (eFluor780 or eFluor450; eBioscience; Cat # 65-0865 or 65-0863). Data were acquired on a CytoFLEX S flow cytometer (Beckman Coulter) and CytExpert(Beckman Coulter software. Prior to data collection, quality controls metrics were checked using quality control beads (CytoFLEX Daily QC Fluorospheres; Beckman Coulter; B53230). Data were compensated using single positive stained cells. Fluorescence minus one (FMO) control stained cells were used for setting gates for analysis. Data were analyzed using FlowJo™ Software (BD Life Sciences). The FlowJo Proliferation Platform was used to calculate proliferation and division index. Proliferation index was calculated as the total number of divisions divided by the number of cells that went into each division. Division index was calculated as the average number of cell divisions that a cell from the original donor population underwent.

### Primary rat cortical neuron-glia co-cultures

Primary cortical neuron-glia co-cultures were prepared from neonatal Sprague-Dawley rats postnatal day 0-1. Pup sex was determined by measuring the anogenital distance, and number of male and female pups determined to equalize male/female ratio. The pups were euthanized by decapitation and their brains excised, and the neocortex dissected in ice-cold Hanks’ Balanced Salt Solution (Gibco, Thermo Fisher Scientific, Waltham, MA, USA; Cat #14185-052) supplemented with 1 M HEPES buffer (pH 7.55; Sigma-Aldrich, St. Louis, MO, USA; Cat #BP310-500) and pooled together in Hibernate A (Gibco, Thermo Fisher Scientific; Cat #A1247501). Tissue was then cut into smaller chunks using sterile razorblades and then incubated at 37°C for 23 min in Hibernate A (Gibco, Thermo Fisher Scientific; Cat #A1247501) containing 2.3 mg/mL papain (Worthington, Lakewood, NJ, USA; Cat #LS003119) and 95 μg/mL DNase (Sigma-Aldrich; Cat #D5025). At the end of the incubation, papain/DNase solution was removed, and the tissue rinsed with Neurobasal Plus medium (Thermo Fisher Scientific; Cat #A3582901) supplemented with 2% B27 (Thermo Fisher Scientific; Cat #A3582801), 1% GlutaMAX (Thermo Fisher Scientific; Cat #35050-061), 10% horse serum (Gibco, Thermo Fisher Scientific; Cat #26050-088) and 1 M HEPES buffer. Papain-digested tissue was then physically triturated using bent tips pipettes (Bellco, Vineland, NJ, USA; Cat #:1273-40004). Cells were counted using a Cellometer Auto T4 Automated Cell Counter (Nexcelom Bioscience LLC, MA, USA) and plated on 6 well-plates (Nunclon Delta Surface, Thermo Fisher Scientific; Cat #140675) precoated with 500 μg/mL poly-L-lysine (Sigma-Aldrich; Cat #P1399). Cells were seeded at 100.000 cells/ cm^2^ on each well and allowed to settle and attach at 37°C under 5% CO_2_. Three to four hours post-plating, culture medium was replaced with growth medium (Neurobasal Plus basal medium supplemented with 2% B27 and 1% GlutaMAX). On day *in vitro* (DIV) 4, half of the conditioned medium was replaced by fresh growth medium supplemented with 10 µM cytosine-arabinoside (AraC) at a final concentration of 5 µM AraC to inhibit glial proliferation.

### Cortical neuron-glia co-cultures stimulation

At DIV 9, growth medium was removed, and cells were treated with 100 nM of 11-Cl-BBQ in fresh growth medium or with the equivalent volume of growth medium containing just the vehicle (0.1% DMSO). Each treatment was conducted in triplicate. At DIV 12, after incubating at 37ºC for 3 days, cells were detached from the plate using a cell scraper, transferred to a 1.5ml microtube and centrifuged at 300 x g for 3 min. The pellet was washed with 1X DPBS (Dulbecco’s phosphate buffered saline, GIBCO, Cat #14190-144), and recentrifuged. The supernatant was discarded and pelleted cells were flash-frozen with liquid nitrogen for RNA extraction.

### Keratinocyte culture

A minimally deviated line of spontaneously immortalized human keratinocytes (23) were grown with 3T3 feeder layer support in a 2:1 mixture of Dulbecco’s Modified Eagle Medium (DMEM; Thermofisher; Cat # 12100061) and Ham’s F-12 nutrient mix media (Thermo Fisher; Cat # 21700075) supplemented with 5% fetal bovine serum (R&D systems; cat# S11150), 0.4 μg/ml hydrocortisone (Sigma; Cat # 3867), and 10 ng/ml epidermal growth factor (GenScript; cat # Z00333). As keratinocytes attached more tightly to the dishes, they outcompete 3T3 fibroblast cells when reaching confluence. At confluence, keratinocytes were treated with 100 nM 11-Cl-BBQ in medium without epidermal growth factor for 24 hours before harvested for analysis.

### Real-time PCR

RNA isolation was performed using the RNeasy mini kit (Qiagen; Cat # 74104). cDNA was synthesized using the High-Capacity cDNA Reverse Transcription Kit (Applied Biosystems; Cat # 4368814). qPCR reactions were performed using SYBR™ Green PCR Master Mix (Applied Biosystems; Cat # 4309155). Nrp1, Ahr, and Cyp1a1 levels were normalized to Actb using primers from Integrated DNA Technologies.

*Actb* Forward: AATCGTGCGTGACATCAA Reverse: GCCATCTCCTGCTCGAAG

*Nrp1* Forward: ATAGCGGATGGAAAACCCTGC Reverse: GGCTGCCGTTGCTGTGCGCCA

*Ahr* Forward: GGAAAGCCCGGCCTC Reverse: CTGGTATCCTGTTCCTGAATGAATTT

*Cyp1a1* Forward: GGTGGCTGTTCTGTGAT Reverse: AAGTAGGAGGCAGGCACA

### Statistics

All statistical analyses were performed using Prism (Graphpad; v9.5.1; https://www.graphpad.com/). An unpaired Students *t*-test was performed to compare the mean between two treatment groups. One-way ANOVA with Tukey’s test for multiple comparisons was used to determine differences between treatment groups. A two-way ANOVA with Bonferroni’s test for multiple comparisons was used to compare two variables. p ≤ .05 was considered statistically significant.

## 3 Results

### 3.1 AhR activation prevented the increased Nrp1 expression on proliferating CD4^+^Foxp3^-^ cells *in vivo*

Based on previous studies demonstrating that Nrp1 expression on Foxp3^-^ cells correlated with inflammatory disease severity(18, 19), we hypothesized that Nrp1^+^Foxp3^-^ T cells would increase over the course of CD4^+^ T cell activation. Using a parent-into-F1 alloresponse model of T cell activation, donor CD4^+^ cells were analyzed at different time points following adoptive transfer. Consistent with the CD4^+^ T cell dependent stage of the alloresponse (19), there was a significant increase from ∼15% to 65% of CD4^+^ T cells that were Nrp1^+^Foxp3^-^ over the first four days (Figure 1A). The percentage of Nrp1^+^Foxp3^-^ cells continued to increase until day 15, albeit at a decreased rate in comparison to the initial four days of the response. We previously showed that AhR activation prevented the development of the alloresponse by inducing Tr1 cell differentiation and inhibiting the development of cytotoxic T lymphocytes, host cell death, and associated weight loss (19). Prevention of the alloresponse by the high affinity AhR ligands, TCDD and 11-Cl-BBQ, was dependent on AhR signaling in CD4^+^ T cells within the first 3 days post-adoptive transfer; AhR ligands were unable to prevent the clinical manifestations of this graft-versus-host response when T cells were transferred from AhR^−/−^ mice (20, 24). Consistent with those findings, TCDD and 11-Cl-BBQ, significantly reduced Nrp1^+^Foxp3^-^ CD4^+^ T cells throughout the alloresponse (Figure 1A).

**Figure 1:**
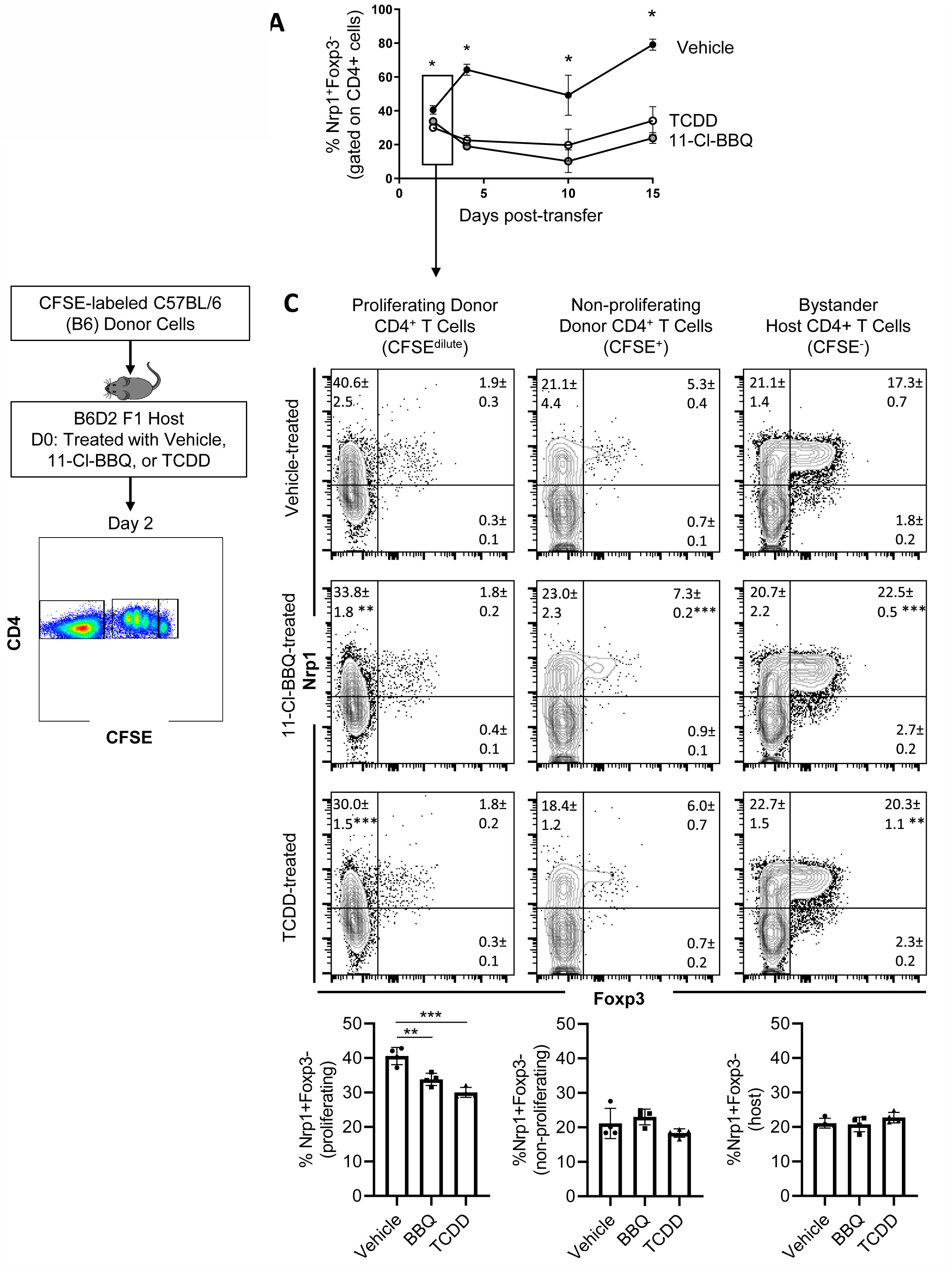
Nrp1 expression increases on CD4^+^Foxp3-cells following activation in vivo, and this increase is prevented when AhR is activated. A. CFSE-labeled C57BL/6 (B6) donor cells were transferred into the B6D2F1 host. The percentage of Nrp1^+^ FoxP3-CD4^+^ cells were measured over 15 days in vehicle, TCDD, and 10-Cl-BBQ-treated mice. B. Schematic of gating strategy for day 2 analysis. C) Flow cytometry plots of representative mice. Nrp1 and Foxp3 were stained in alloresponsive donor CD4^+^ cells (CFSE^dilute^), non-responding CD4^+^ T cells (CFSE^hi^), and bystander host CD4^+^ T cells (CFSE^-^). n=4 mice/group representing one of two independent experiments. Each data point represents an individual mouse. A one-way ANOVA was performed. ^*^ indicates p-value <0.05. ^**^ indicates p-value <0.01. ^***^ indicates p-value <0.001. Numbers in the quadrant represent the mean and standard deviation of the percentage of cells in the gate.

Newly activated CD4^+^ T cells are particularly sensitive to AhR ligands, with AhR-mediated transcriptional and phenotypic changes, including the downregulation of Nrp1, occurring on day 2 (9). To determine how AhR activation impacted proliferating, non-proliferating, and bystander host CD4^+^ T cells, donor cells were labeled with CFSE prior to adoptive transfer (Figure 1B). On day 2, proliferating (CFSE dilute) donor CD4^+^ T cells had a higher proportion of Nrp1^+^Foxp3^-^ cells compared to non-responding (CFSE^hi^), or bystander (host, CFSE^-^) CD4^+^ T cells. That AhR activation only modulated Nrp1 expression in proliferating CD4^+^ T cells suggests that AhR ligands prevent the upregulation of Nrp1 expression rather than inhibiting basal expression (Figure 1C). While Foxp3^+^ cells only made up a small percentage of donor CD4^+^ T cells which limits our analysis of donor cells, the majority of host Foxp3^+^ cells coexpressed Nrp1, consistent with the abundance of data demonstrating that Nrp1 can be a Treg marker (15). On these Tregs, AhR activation did inhibit Nrp1 expression. Instead, Nrp1 was slightly, but significantly, increased on Foxp3^+^ cells following AhR activation.

### 3.2 Nrp1^+^ cells are proliferative and express an effector phenotype

For CD4^+^ T cells to provide help for CTL differentiation, they must first undergo extensive proliferation. Since Nrp1^+^Foxp3^-^ T cells increased four-fold during the first four days of activation, we analyzed CFSE dilution to determine if proliferation of Nrp1^+^ cells was the primary driver of the expanding donor CD4^+^ T cells. On both days 2 and 4, Nrp1^+^Foxp3^-^ splenocytes had undergone more divisions compared to the Nrp1^-^Foxp3^-^ cells, corresponding with a larger proliferation and division index (Figure 2A, 2B). An increased proliferative capacity is consistent with previous findings that *in vitro* activated CD4^+^Foxp3^-^Nrp1^+^ cells have a higher level of the cell proliferation marker Ki67 (21). In the same study, Nrp1^+^Foxp3^-^ cells were found to have an activated phenotype expressing high levels of CD44. To determine if AhR signaling altered the activation status of Nrp1^+^Foxp3^-^ cells, coexpression of CD44 and CD45RB was assessed in donor CD4^+^ T cells isolated from mice treated with vehicle or TCDD (Figure 3A, 3B). The majority (∼90%) of Nrp1^+^Foxp3^-^ cells expressed CD44^hi^CD45RB^lo^, a phenotype consistent with activated cells; this percentage of activated cells was higher than in Nrp1^+^Foxp3^+^, as well as Nrp1^-^ cells (Figure 3C). Interestingly, AhR signaling did not reduce the percentage of Nrp1^+^Foxp3^-^ cells that expressed an activated phenotype, consistent with their ability to proliferate (Figure 2). Instead, AhR activation significantly reduced the total number of Nrp1^+^Foxp3^-^CD44^hi^CD45RB^lo^ cells (Figure 3D). Conversely, TCDD treatment increased the percentage and number of Nrp1^-^Foxp3^-^ cells that express an activated phenotype. This population of cells likely include activated Tr1 cells that we have previously shown are Foxp3^-^. These Tr1 cells are migratory, although, in the current study, we have limited our analysis to the spleen, where we have established that Nrp1^+^Foxp3^-^ cells correlate with the clinical manifestations of the alloresponse(9, 19). Collectively, the data demonstrate that AhR ligands can reduce the overall number, but not the percentage of Nrp1^+^Foxp3^-^ cells that express an activated phenotype.

**Figure 2:**
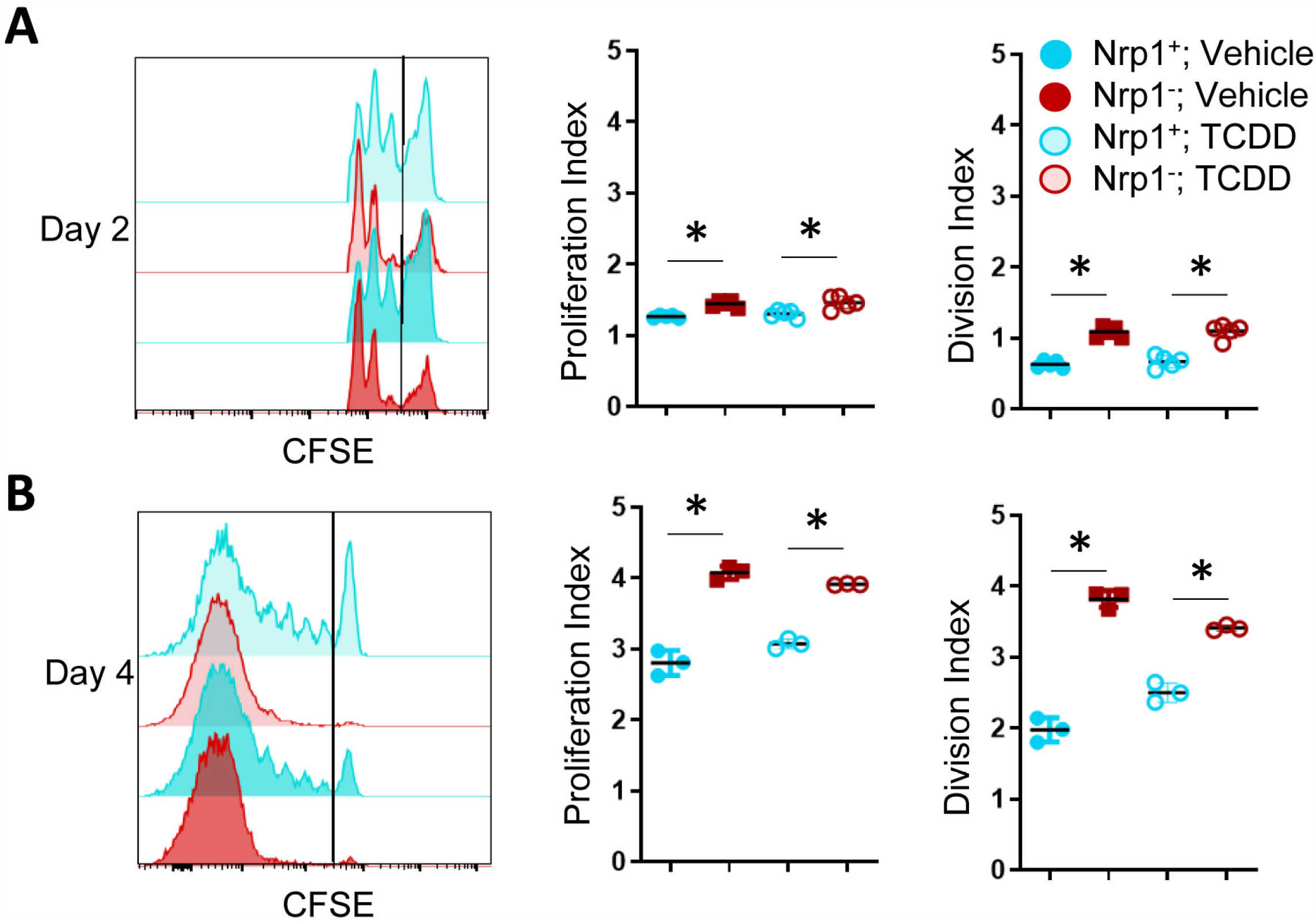
Nrp1^+^ cells have an increased proliferation index and increased division index compared to Nrp1-cells. B6 donor cells were transferred into the B6D2F1 host and treated with vehicle or 15μg/kg TCDD. CFSE-labeled splenocytes were examined on days 2 (A) and 4 (B) of the alloresponse. The proliferation index and division index were examined in CD4^+^ Foxp3-Nrp1^+^ cells and CD4^+^Foxp3^-^Nrp1^+^ donor (HD2^d−^) cells. A one-way ANOVA was performed. ^*^ indicates p-value <0.05 n=3-5 mice/group from two separate experiments. Each data point represents an individual mouse.

**Figure 3:**
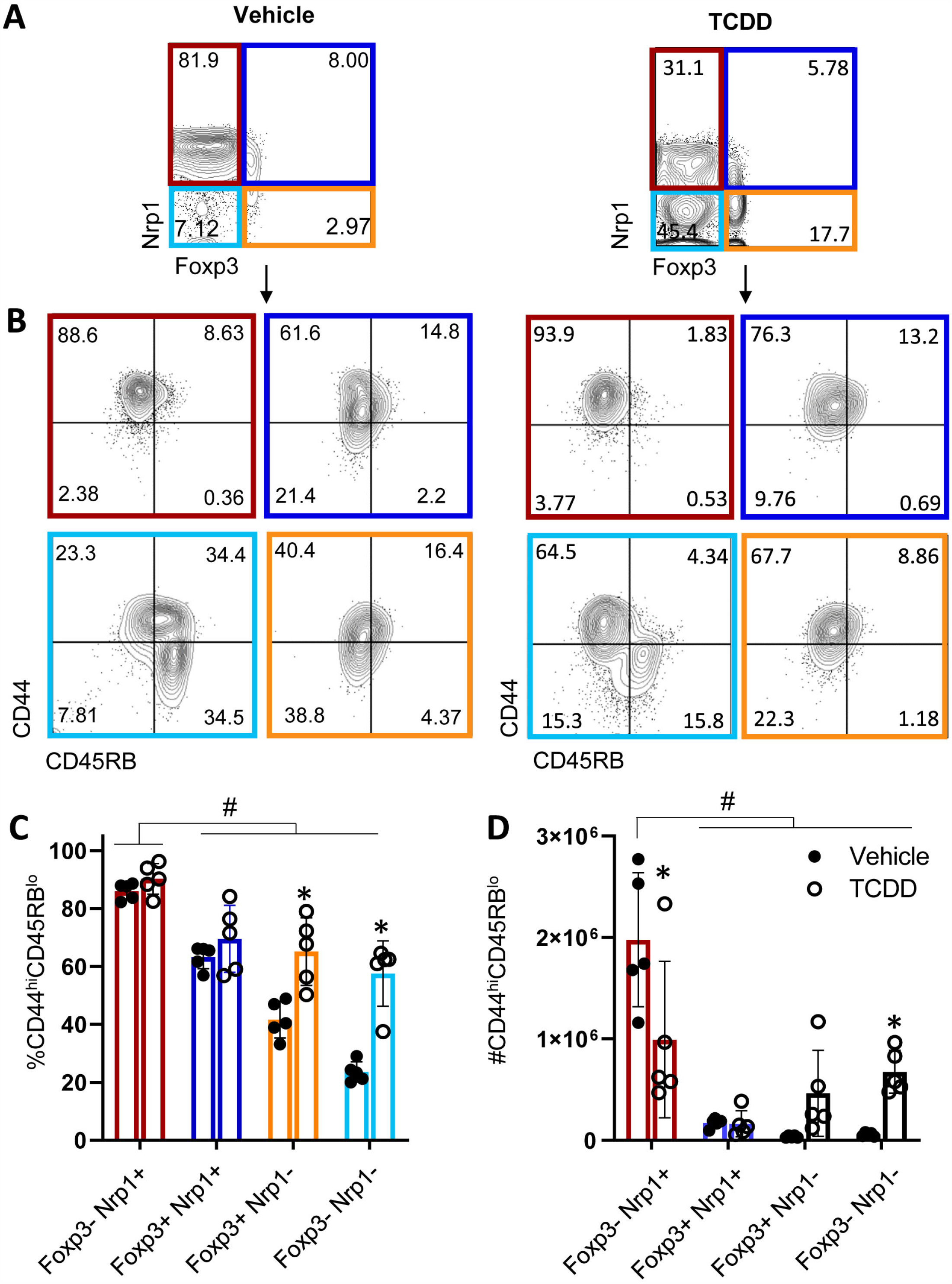
CD4^+^Nrp1^+^Foxp3^-^ cells express an effector phenotype. B6 donor cells were transferred into the B6D2F1 host and treated with vehicle or 15μg/kg TCDD. Following 15 days, splenocytes were stained for H2D^d^, CD4, Foxp3, Nrp1, CD44, and CD45RB. A. Flow cytometry plots of Nrp1 and Foxp3 expression on donor (HD2^d−^) CD4^+^ T cells in vehicle-treated and TCDD-treated mice. B. Flow cytometry plots of CD44 and CD45RB stained cells gated on Nrp1 and Foxp3 expression, shown in A. C. The percentage of CD44hiCD45RBlo cells in gates based on coexpression of Foxp3 and Nrp1 isolated from vehicle and TCDD-treated mice. A two-way ANOVA was performed. ^*^ indicates p-value <0.05. D. The number of CD44hiCD45RBlo cells in gates based on coexpression of Foxp3 and Nrp1 isolated from vehicle and TCDD-treated mice. A two-way ANOVA was performed. ^*^ indicates p-value <0.05 when compared between treatments. # indicates p-value <0.05 when compared between populations. n=5 mice/group from one experiment. Each data point represents an individual mouse.

### 3.3 AhR activation does not globally inhibit CD4^+^ T cell activation, and Nrp1 is a delayed activation marker

Following productive T cell receptor (TCR) engagement, CD4^+^ T cells upregulate a set of activation markers. First among these is CD69 which is regarded as an early activation marker, followed by the IL-2 receptor, CD25 (25). CD29 makes up part of the very late antigen (VLA) complex that is involved in T cell emigration, and, like Nrp1, is among the most downregulated genes in CD4^+^ T cells following AhR activation (9). As the Nrp1^+^Foxp3^-^ cells express elevated proliferation and an increase in cells expressing an activated phenotype (21), Nrp1 was assessed as an activation marker. Following T cell activation with CD3/CD28 beads, the expression of CD29, CD69, and CD25 was upregulated one day following stimulation (21), whereas Nrp1 lagged by one day (Figure 4A).

**Figure 4:**
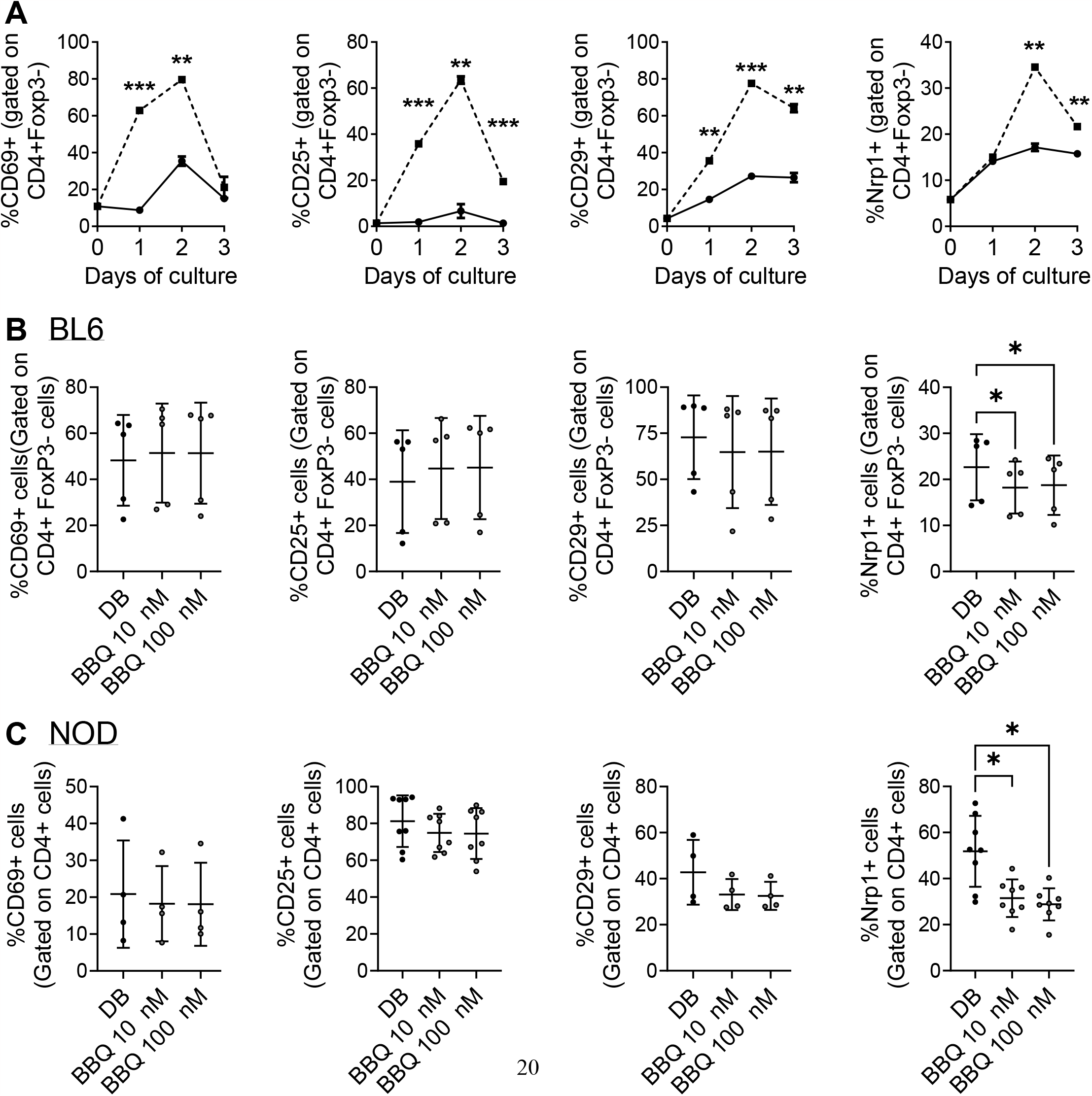
AhR activation does not globally inhibit CD4^+^ T cell activation markers and Nrp1 is a delayed activation marker. A. CD4^+^ cells were cultured unstimulated (—) or with anti-CD3/CD28 Dynabeads (---) over the course of 3 days. The percentage of Nrp1^+^, CD29^+^, CD69^+^, and CD25^+^ cells were gated on CD4^+^ Foxp3^-^ cells. An unpaired t-test was performed. * indicates p-value <0.05. B,C. CD4^+^ cells isolated from C57BL/6 mice(B) and NOD mice (C) were cultured with 1nM Cl-BBQ, 10 nM Cl-BBQ, or 100 nM Cl-BBQ for 2 days. All cells were also co-cultured with anti-CD3/CD28 Dynabeads. Following four days of incubation, cells were stained for Nrp1, CD29, CD69, and CD25. A one-way ANOVA was performed. ^*^ indicates p value <0.05. n=4-8 mice/cell culture condition. Each data point represents cells from an individual mouse and includes at least two independent experiments.

Based on the profound suppression of the alloresponse, it is possible that AhR ligands could simply act by preventing the activation of CD4^+^ T cells. However, AhR ligands have been linked to an upregulation of activation markers (e.g. CD25, CD69, GITR, ICOS) as well as the downregulation of CD62L (9, 19, 24). To determine how AhR signaling influences Nrp1 and other T cell activation markers, CD4^+^ T cells were isolated from C57BL/6 and NOD mice, two strains in which we have observed AhR activation reduces Nrp1^+^Foxp3^-^ cells (18, 19). The expression of CD69, CD25, and CD29 was not altered by increasing concentrations of 11-Cl-BBQ (Figure 4B, Figure 4C). However, AhR activation reduced the expression of Nrp1, consistent with the *in vivo* studies in Figure 1. Of note, AhR is expressed in CD4^+^ T cells by 24 hours in culture (26). Collectively, these data demonstrate that AhR activation does not globally inhibit the expression of CD4^+^ T cell activation markers, but rather preferentially modulates the expression of some CD4^+^ T cell surface proteins.

### 3.4 AhR activation does not universally inhibit Nrp1 upregulation in CD4^+^ T cell subsets

AhR ligands prevented the upregulation of Nrp1 expression on CD4+T cells stimulated with anti-CD3/CD28 (Figure 4B and C). We next set out to evaluate whether this pattern holds true following differentiation into T helper subsets. We previously found that AhR activation did not alter the ability of CD4^+^ T cells to differentiate into helper subsets *in vitro*, although CD4^+^ T cells from AhR^−/−^ mice had increased *Il17a* expression regardless of polarizing condition (26). In the current study, CD4^+^ T cells were isolated from prediabetic NOD mice and polarized to Th0, Th1, Th2, Th17, and Treg subsets, and Nrp1 expression was assessed following AhR activation. CD4^+^ T cells from NOD mice were used for these studies since the reduction in Nrp1 expression was more pronounced in the *in vitro* studies using CD4^+^ T cells from this strain compared to C57BL/6 mice (Figure 4C). Nrp1 expression was analyzed on CD4^+^Foxp3^-^ cells, although Foxp3^+^ cells were not initially removed prior to polarization. 11-Cl-BBQ reduced the percentage of Nrp1^+^ cells under Th1-polarized conditions, whereas Nrp1 expression was not reduced on CD4+ T cells polarized under the other conditions (Figure 5A). Confirming that AhR activation does not simply inhibit all T cell activation markers, the proportion of CD25^+^ CD4^+^ cells was unaltered (Figure 5B). However, when Nrp1 expression was examined on CD25^+^ gated cells, there was a significant reduction in the percentage of Nrp1^+^ cells in both Th1 and Treg cells, and a trend toward a reduction in Th0 and Th17 cells. In contrast, Nrp1 expression was unaltered in Th2-polarized cells (Figure 5C). This pattern was dependent on AhR expression, as 11-Cl-BBQ did not alter Nrp1 expression on CD4^+^ T cells from AhR^−/−^ NOD mice. (Figure 5D-F). Given that 11-Cl-BBQ did not downregulate Nrp1 expression in the Th2 CD4^+^ T cell subset, the gene expression of *Ahr* was assessed to determine if this subset was unresponsive because the target was not expressed. Indeed, Th2 cells had comparatively low expression levels of *Ahr* (Figure 5G). Consistently, the upregulation of *Cyp1a1* was not as pronounced in Th2 cells as it was with the other T helper subsets in the presence of 11-Cl-BBQ (Figure 5H).

**Figure 5:**
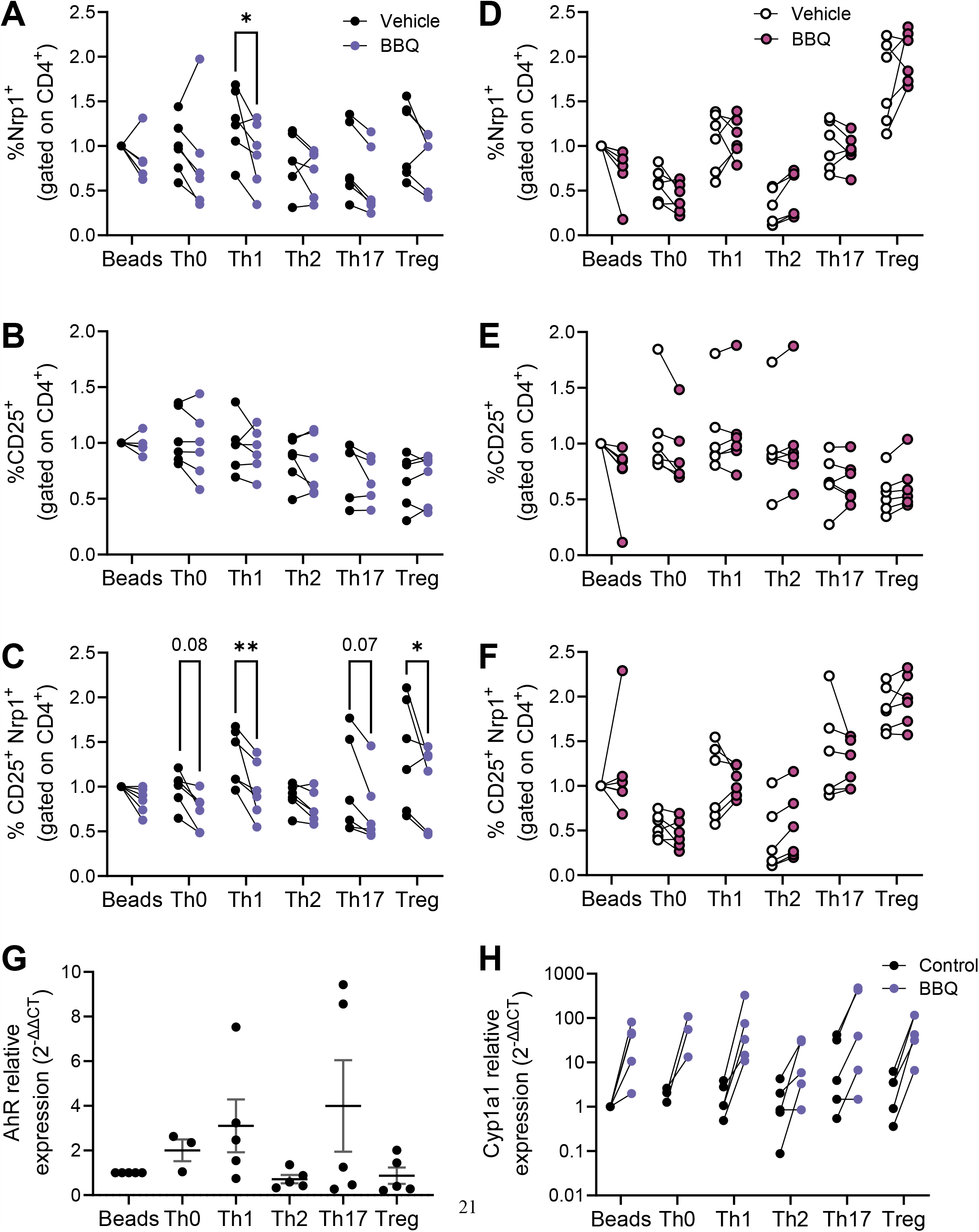
AhR activation downregulates Nrp1 expression in Th1 and Treg subsets. CD4^+^ T cells isolated from the spleen of NOD mice were cultured under conditions polarized towards Th0, Th1, Th2, Th17, or Treg. All cells were also co-cultured with CD3/CD28 Dynabeads. AhR was activated with 100 nM Cl-BBQ. Cells were incubated at 37°C for 4 days. Cells from WT mice were stained for Nrp1, CD25, and CD4. The percentage of Nrp1^+^ (A), CD25^+^ (B), and Nrp1^+^CD25^+^ (C) were examined. Cells from AhR KO mice were stained for Nrp1, CD25, and CD4. The percentage of Nrp1^+^ (D), CD25^+^ (E), and Nrp1^+^CD25^+^ (F) were examined. qPCR analysis was performed on cells from WT mice. The relative expression of *Ahr* (G), and *Cyp1a1* (H) is shown. Data were normalized to stimulated control. For A-F, a two-way AVOVA was performed. ^*^ indicates p-value <0.05. ^**^ indicates p-value <0.01. n=5-6 mice/cell culture condition. Each data point represents cells from an individual mouse and includes at least two independent experiments.

### 3.5 AhR-mediated inhibition of Nrp1^+^ expression is dependent on IL-2

AhR and Nrp1 are both expressed in several cell types including neurons, keratinocytes, and several cancer cell lines(11, 27-31). To determine if AhR activation will decrease Nrp1 expression in other cell types or if the downregulation of Nrp1 occurs solely in CD4^+^ T cells, Nrp1 expression was measured in primary rat cortical neuron-glia co-cultures, and keratinocytes following co-culture with 11-Cl-BBQ. In contrast to CD4^+^ T cells, AhR activation upregulated Nrp1 expression in neuron-glia cultures (Figure 6A) and did not alter Nrp1 expression in keratinocytes (Figure 6B). Conversely, there is an upregulation in Nrp1 expression following AhR activation in neuron-glia co-cultures. These data suggest that AhR signaling alone does not directly inhibit Nrp1 expression.

**Figure 6:**
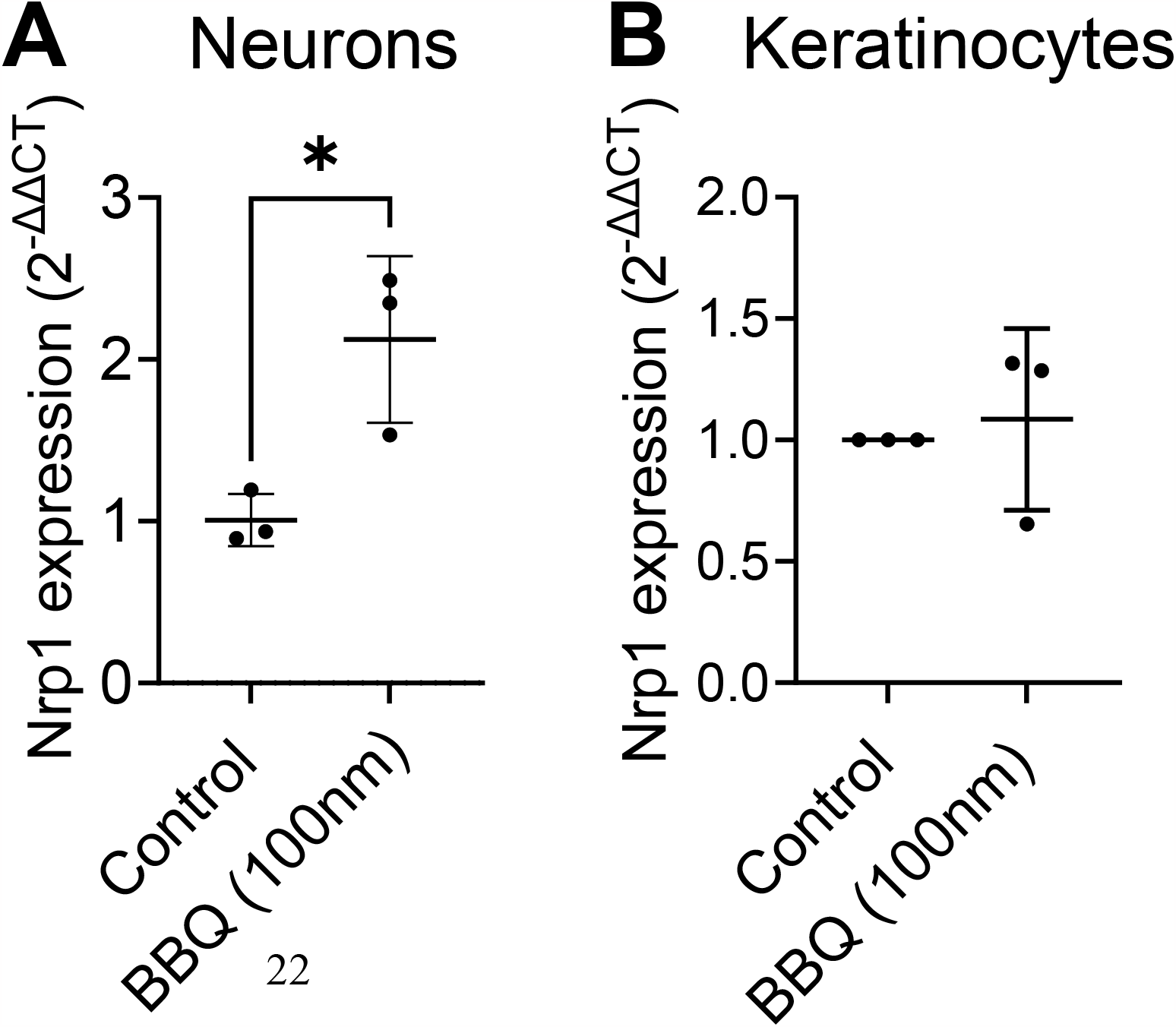
AhR ligands do not downregulate Nrp1 expression in primary rat-cortical neuron-glial co-cultures or keratinocytes. Cortical neurons and keratinocytes were treated with 100nM 11-Cl-BBQ (, and qPCR analysis was performed. The relative expression of Nrp1 is shown. An unpaired t-test was performed. ^*^ indicates p-value <0.05. n=3 cell culture wells/condition.

Given that activated CD4^+^CD25^+^T cells, but not non-proliferating T cells, keratinocytes, or neurons, were most sensitive to AhR-mediated downregulation of Nrp1 and that CD25 is the receptor for T cell growth factor IL-2, the inhibition of Nrp1 upregulation was hypothesized to be dependent on IL-2. To determine whether AhR-IL-2 crosstalk is required to prevent Nrp1 upregulation, IL-2 was neutralized *in vitro* and *in vivo*. Since IL-2 is necessary for CD4+ T cell proliferation *in vitro*, and early production of IL-2 is a required initiating event in the parent-into-F1 alloresponse (32), antibody concentrations were selected to neutralize excess IL-2 production (9) without impairing baseline T cell proliferation and Nrp1 upregulation. *In vitro*, increasing concentrations of anti-IL-2 reversed the reduction Nrp1 expression following AhR activation (Figure 7A). Similarly, when IL-2 was neutralized *in vivo*, TCDD was unable to reduce Nrp1 expression on alloresponding CD4^+^CD25^+^ T cells after treatment with TCDD (Figure 7B). This reversal is consistent with our previous findings using the alloresponse model that an increase in IL-2 occurs early after AhR activation, and is necessary for the immunoregulation of newly activated CD4^+^ T cells (9). These data suggest that the mechanism by which AhR acts to inhibit the upregulation of Nrp1 is dependent on the presence of IL-2.

**Figure 7:**
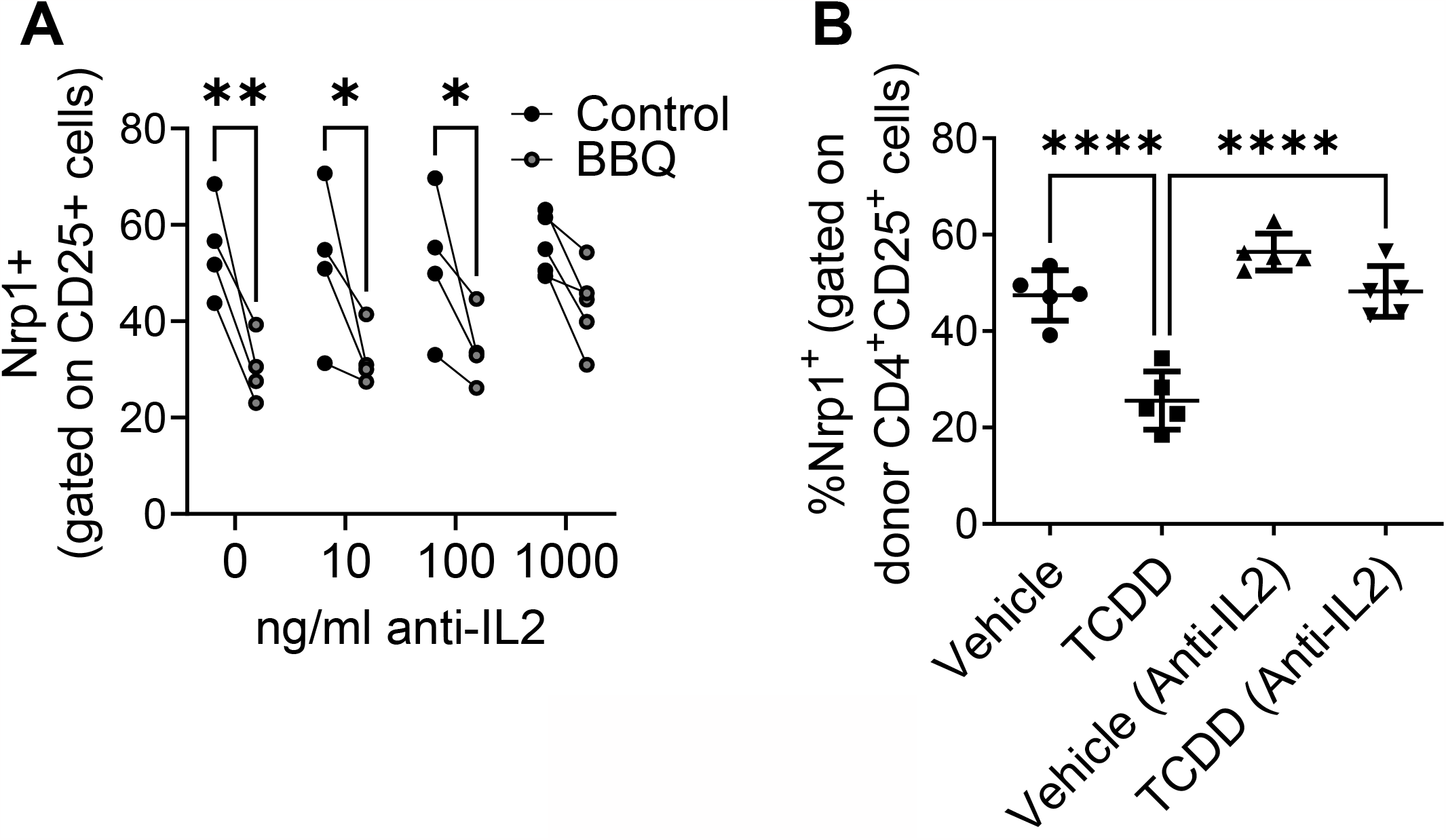
The reduction of CD25^+^ Nrp1^+^ expression resulting from AhR activation is dependent on IL-2. A. CD4^+^ T cells isolated from NOD mice splenocytes were co-cultured with 1000 ng/ml, 100 ng/ml, 10 ng/ml or 0 ng/ml of anti-IL-2. All cells were also co-cultured with IL-12, anti-IL4, and CD3/CD28 Dynabeads. AhR was activated with 100 nM Cl-BBQ. Following incubation at 37°C for 4 days, cells were stained, and Nrp1 expression was analyzed on CD4^+^CD25^+^Foxp3^-^ cells. A two-way ANOVA was performed. ^*^ indicates p-value <0.05. ^**^ indicates p-value <0.01. n=4 mice/condition. B. CFSE-labeled cells from B6 mice were injected into F1 hosts and treated i.p. with anti-IL-2 or isotype control and 15μg/kg TCDD or vehicle. Nrp1 expression on proliferating donor (CFSE dilute) CD4^+^CD25^+^ cells were analyzed on day 2.A one-way ANOVA was performed. ^***^ indicates p-value <0.001. n=5 mice/group. Each data point represents an individual mouse.

## 4 Discussion

AhR activation by high affinity ligands promotes Treg differentiation, which is accompanied by suppression of proinflammatory CD4^+^ T cell subsets. The molecular sequelae leading to Treg differentiation have been well studied; however, it is less clear if AhR signaling concurrently inhibits the expression of genes involved in effector T cell function. In the current study, we found that AhR activation directly alters CD4^+^ T cell activity by inhibiting the upregulation of Nrp1 on CD4^+^Foxp3^-^ cells. Inhibition of Nrp1 expression was most pronounced in Th1 cells and was dependent on IL-2 signaling. These results suggest that Nrp1 regulation could be a mechanism by which AhR ligands lead to immune suppression of CD4^+^ T cell responses.

Although primarily used as a Treg marker (15), several studies have suggested that Nrp1 is also involved in CD4^+^ effector T cell activation. FACS data shows Nrp1 is expressed on CD4^+^ Foxp3^-^ cells in addition to Foxp3+ cells (18, 19, 21, 33-35), and these cells can increase during immune-mediated diseases. In NOD mice, Nrp1 expression in CD4^+^Foxp3^-^ cells exhibited a strong correlation to islet infiltration (18). Similarly, we previously showed that CD4^+^Nrp1^+^Foxp3^-^ cells have a positive correlation with cytotoxic T lymphocytes in an alloresponse model (19). In the current study, we found that alloresponsive Nrp1^+^Foxp3^-^ cells expressed an activated phenotype. This CD44^hi^CD45RB^lo^ phenotype is consistent with Nrp1^+^ conventional T cells found under inflammatory conditions in TGF-β RII-deficient mice (17) (Weiss 2012), and, more recently, in a TCR transgenic model of diabetes (21). Similarly, the authors found that Nrp1 expression was induced following *in vitro* stimulation of CD4^+^Foxp3^-^, with cells expressing Nrp1 having increased expression of IFNγ and TNFα compared to Nrp1^-^Foxp3^-^ cells. These cells had increased proliferative potential, consistent with the findings of our current study (21).

The current study focused on Nrp1 expression on host alloresponding CD4^+^ T cells; however, there is concurrently an expansion of host CD4^+^ T cells over the first 4 days in the acute parent-into-F1 alloresponse model preceding the elimination phase that begins on day 10 (36). This expansion is likely a result of antigen-independent “bystander” CD4^+^ T cell activation, which is more commonly observed in memory CD4^+^ T cells in response to cytokines (e.g. STAT inducers like IL-2) or microbial/TLR stimulation (37). That AhR signaling does not interfere with Nrp1 expression on these bystander-activated CD4^+^ T cells in the same manner as alloantigen-activated naïve CD4^+^ T cells is consistent with previous findings that newly activated CD4^+^ T cells are more sensitive to AhR ligands. The sensitivity of CD4^+^ T cells to differentiation following AhR signaling diminishes 3 days after activation (38), and fully differentiated effector/memory T cells are not significantly affected by AhR ligands (38, 39). Furthermore, single cell RNAseq data demonstrated that memory cells stimulated with Th1 cytokines or type 1 interferon do not express *Ahr* (40). Furthermore, bystander activation of CD4^+^ T cells by IL-2 tends to promote Th2 differentiation, which we found least susceptible to Nrp1 downregulation (41).

We used two complementary models to interrogate the impact of AhR signaling on Nrp1 expression on CD4+ T cells. However, both the parent-into-F1 alloresponse and the NOD mouse have a Th1 bias. We found that *in vitro* polarized Th1 cells from NOD mice were most sensitive to AhR-dependent downregulation of Nrp1 expression, with Th2 cells the least sensitive to both AhR signaling and Nrp1 modulation. In these studies, CD4^+^ T cells were isolated from splenocytes of prediabetic adult NOD mice; although they were not yet hyperglycemic, an underlying Th1-skewed autoimmune response would have already been ongoing in these mice. Further studies would be needed to see if downregulation of Nrp1 on effector cells similarly occurs in Th2-biased mouse models (e.g. BALB/c); Abberger, et al previously showed that Nrp1 expression is upregulated on activated CD4^+^Foxp3^-^ cells from BALB/c mice (21).While the inhibition of Nrp1 expression on newly activated CD4^+^ T cells following AhR activation is evident, the mechanism by which Nrp1 expression is regulated is less clear. As a transcription factor, AhR has been implicated in both the up- and downregulation of gene expression; however, our data show that AhR does not inhibit the expression of Nrp1 across all cell types. Specifically, Nrp1 expression is not altered by AhR ligands in keratinocytes or in Th2 cells and is surprisingly increased in neurons. These data suggest that AhR alone does not directly downregulate Nrp1. Instead, AhR may require additional binding partners expressed under specific conditions or act indirectly through the induction or repression of Nrp1 transcriptional regulators. In studies interrogating Nrp1 expression in different cell types, both transcriptional activators and repressors of Nrp1 have been reported that have crosstalk with the AhR pathway. In pancreatic neuroendocrine neoplasms, the IL6/Stat3 pathway upregulates Nrp1 (42), and AhR signaling has been shown to inhibit IL-6 (43-45). In the mammary glands of MMTV-Wnt1 mice, Nrp1 is upregulated by Wnt/b-catenin signaling (13) and AhR negatively regulates this pathway during early stem cell differentiation (46). In the current study, we found that the AhR-mediated downregulation of Nrp1 is dependent on IL-2 *in vivo* and *in vitro*. One possible crossover pathway between AhR, Nrp1, and IL-2 is through E2F7 and E2F1. E2F7-HIFα complex is thought to be a repressor of Nrp1 through inhibiting E2F1, an activating protein that binds directly to the Nrp1 promoter (47, 48). Although we previously reported global gene expression analysis of alloresponding donor CD4^+^ T cells isolated from AhR ligand-treated mice, it has become clear that this donor population encompasses a heterogenous mix of proinflammatory effector T cells and Tr1 cells. In the future, single cell RNAseq on activated CD4^+^ T cells may help uncover coregulatory pathways between AhR and Nrp1.

Because AhR activation leads to the downregulation of Nrp1 on effector T cells while still promoting Tregs, it can be considered an attractive therapeutic target for a range of immune-mediated diseases. Furthermore, Nrp1 might have utility to serve as a useful biomarker for tracking the development of inflammatory diseases or the efficacy of treatment. Collectively, our data highlight AhR-IL2-Nrp1 signaling as a novel mechanism by which AhR modulates effector CD4^+^ T cell responses.

## 6 Author Contributions

SS, KM, KT, FDCS, LWL, JP, and AE performed experiments. SS, KM, MG, and AE analyzed the data and generated figures. AE conceived the study. KM, NK, and AE provided funding for the project. SS, KM, CM, LW, FDCS, PL, and AE wrote the manuscript. All authors contributed to manuscript revision, read, and approved the submitted version.

## 8 Acknowledgments

The authors would like to thank the UC Davis Mouse Biology Program; Heather Kahalehili, Jack Connolly, Madelynn Tucker, and Jocelyn Molina for genotyping and laboratory support; Robert Rice and Lisa Miller for helpful discussions.

